# Evaluation of entomopathogenic nematodes against red palm weevil, *Rhynchophorus ferrugineus* (Olivier) (Coleoptera: Curculionidae)

**DOI:** 10.1101/2022.03.02.482635

**Authors:** Gul Rehman, Muhammad Mamoon-ur-Rashid, Atiq Ahmad Alizai

## Abstract

The entomopathogenic nematodes play a pivotal role as bio-control agents of different species of insect pests including red palm weevil. In current investigations, infective capabilities of four species of entomopathogenic nematodes including *Hetrerorhabditis bacteriophora*, *Steinernema feltiae*, *Steinernema glesri* and *Steinernema carpocapsae* were evaluated against larvae, pupae and adult red palm weevil, under laboratory and field conditions. The pathogenic potential of selected nematode species was assessed based on dissection and adult emergence of weevils. Results indicated that *S. carpocapsae* and *H. bacteriophora* with 94.68 and 92.68% infection were found most effective EPN species against red palm weevil larvae. Focusing the adult emergence, aforementioned EPNs were found most pathogenic when pupae of red palm weevil were treated and resulted into 83.60 and 80.20% infested pupae. It was noted that the adult emergence is the better option for the evaluation of pathogenic potential of EPN compared to dissection of insect. The *S. carpocapsae* was found most effective against 6^th^ instar larvae of red palm weevil and caused 100% mortality at 340 hours after treatment; whereas; *S. glesri* and *S. feltiae* were found least pathogenic and caused 70 and 76% mortality. All the evaluated nematode species were found highly infective under field conditions. The *S. carpocapsae* was found most pathogenic causing 83.60% mortality of red palm weevil. Overall; the tested nematodes were found most effective against larvae followed by adult weevils. The tested nematode species were found least effective against pupae of red palm weevil. Based on current findings, it is concluded that the tested species of nematodes can be used as sustainable option for the management of red palm weevil.

## Introduction

The date palm (*Phoenix dactylifera* L.) belongs to palm family Arecaceae is an important fruit crop and economic resource for many countries. It is the oldest fruit crop cultivated since the pre historic times mostly in the arid regions across the globe having more than 2400 species belonging to 200 genera [19, 34, 38]. Worldwide, over 100 million date palms are grown on an area of 1 million hectare with a total production of about 7.5 million metric tons of date fruit [9]. Date fruits are considered as a complete diet and are a rich source of different nutrients. Dates carry abundant amounts of carbohydrates, dietary fiber, various minerals (Calcium, Iron, magnesium, phosphorus, potassium, zinc, Sulphur, cobalt etc.), proteins and lipids in trace amounts [37]. The date fruit contains 70% carbohydrates, while date proteins are rich in amino acids [39]. The date fruits also have many medicinal values and can play a leading role in preventing deficiency of vital nutrients in developing countries around the globe [25].

A large variety of insect pests attack on various species of palms across the world which has been divided into pre and post-harvest pests. About 112 species of insect pests and mites has been reported to attack date palm including 22 species which attack post-harvest stored dates [8, 27]. Among these insect and mite pests, red palm weevil, *Rhynchophorus ferrugineus* (Olivier 1790) (Coleoptera: Curculionidae) is considered as the most damaging and invasive pests of date palm. So far, it has been documented from 54 countries around the globe, damaging more than 40 palm species from 23 palm genera of which date palm, coconut and the canary island palm are significant [3, 13].

The adult female of red palm weevil lays up to 200 eggs at the base of fresh leaves and in wounds in concealed places of the stem. The neonate grubs bore into the soft fibers and reach the terminal bud tissues reaching a size of about 5 cm before pupation [33]. The larvae feed on soft tender tissues of various date palm species. Just before pupation, the full-grown larvae move towards inner tissues making tunnels and construct cocoons from dried fibers. Larval tunneling encourages the infestation of secondary pests and pathogens (e.g., fungal pathogens) [6, 7, 17]. As the larvae feed on the internal tissues of the palms the initial infestation is extremely difficult to detect. The infestation is usually noticed after the palm tree has been seriously injured [32]. The symptoms of infected palms include tunneling in the tree stem from which yellow color fluid oozes out, chewed up fibers around the holes of tunnels, gnawing sound made by the growing grubs, pupal cases and adults in the leaf axils, fallen pupal cases near the base of the date tree. A very typical sign of red palm weevil’s infestation is distorted crown in case of severe damage which can be easily seen compared to other symptoms of weevil’s infestation [26]. They are usually found infesting any place within the palms from the ground surface of the trunk up to the apical bud [5]. In date palm, about 70% infestation was recorded from the ground up to 1–1.5 m whereas; on *P. canariensis*) the 80–90% infestation was found on the apical portion of the tree [24].

The management of red palm weevil has always been a great challenge for entomologists since the start of modern agriculture throughout the world. Different methods have been employed for the management of Red palm weevil including preventive and curative, bioacoustics detection, chemical detection, thermal imaging detection, host plant resistance, phyto sanitation and agro techniques and integrated pest management program. However, the main thrust of date palm farmers is on the repeated and excessive use of synthetic chemicals. The indiscriminate use of synthetic chemicals causes many problems like development of resistance in insect pests, accumulation of residues on date palm deterioration of environment and causes serious health hazardous to human being [31].

EPNs are a group of soil-dwelling organisms belonging to Steinernematidae and Heterorhabditidae families are well-known natural enemies of insect pests. They play a pivotal role as bio control agents of different insect pests including red palm weevil [15]. The stage of third infective juvenile of the EPN is a non-feeding stage which actively looks for target insect pests by utilizing its host seeking ability. It invades the host insect and enters the insect body through natural openings like spiracles, mouth or anus or thin parts of the cuticle and releases a host-specific bacterium Photorhabdus in case of Heterorhabditidae) and (Xenorhabdus in case of Steinernematidae which kills the target insect pests through bacterial septicemia [10, 16, 40]. The entomopathogenic nematodes are easy to culture and are inexpensive, live for weeks during the infective stages and are capable of infecting a broad range of insect pests including red palm weevil [15,16].

## Materials and Methods

### Red palm weevil culture

The larvae and adults of red palm weevil were collected from date palm trees at Germ Plasm Unit (GPU) date palm orchard in Dera Ismail Khan, Khyber Pakhtunkhwa (KPK), Pakistan. The collected stages were shifted to plastic jars (13 × 3 cm^2^) for mass rearing in the laboratory of Department of Entomology, Gomal University, Dera Ismail Khan. The colonies of red palm weevil were maintained in incubators maintained at constant conditions of 27 ± 2°C, 65 ± 5% R.H. and 14:10 h D:L regimes.

The collected larvae were provided soft portion (30-gram) of date palm (Cv. Dhakki) stems for two days. The jars were washed with tap water after every two days and were completely air dried under sunlight. The mouths of the jars were covered with fine mesh gauze for ventilation purpose. The food of the weevils was changed on every alternate day. The full-grown larvae were provided with a 5-inch sugarcane piece to facilitate cocoon formation. The newly formed pupal cocoons were collected and transferred on daily basis to new jars for adult emergence. The emerged adult weevils were cultured in separate jars having size of 30×60×60 cm^3^ for feeding, mating and oviposition purposes. The adult weevils were provided with pieces of stem along sugar and honey solution to facilitate egg laying. The red palm weevil adults that emerged on the same day were considered of uniform age and were used for the studies. The newly laid eggs were collected on daily basis to ensure supply of age-specific different stages of red palm weevil for the investigations.

### Entomopathogenic nematodes (EPNs)

Entomopathogenic nematodes were purchased through online system from storage bank of nematodes hub Sc Garden Nema Seek™, USA (10 × 10^8^ nematodes). The nematodes were cultured on 5^th^ instar larvae of wax moth (*Galleria mellonella*). The 9 cm plastic Petri dishes were used for the rearing of nematodes and white filter paper was placed in the petri dishes. To maintain moisture level in the petri dishes, 1.5 ml tap water along with ≈500 nematodes using micropipette were added in each petri dish. Ten active larvae of wax moth were introduced in each petri dish using fine camel hair brush with different species of selected nematodes. The petri dishes were sealed using a strip of Para film and petri dishes were placed in an incubator maintained at 18°C for 48 hours. After 48 hours, the petri dishes containing wax moth larvae were removed and dead larvae were separated using white traps for the extraction of infective juveniles following standard method described by Kaya and Stock. The newly emerged infective juveniles were collected on daily basis and stored at 18°C in distilled sterilized water for future use. The EPN of less than 5-day old were used for the investigations.

## Laboratory Studies

### Pathogenicity of EPN’s against Red palm weevil larvae based on dissection

The laboratory experiment was conducted to find out the pathogenicity of four EPNs species against 5^th^ instar larvae of red palm weevil. The experiment was conducted in transparent plastic boxes (9 × 5 × 5cm) lined with Whatman No.1 filter papers. For the studies, 0.1 ml solution containing nematodes (approximately 100 IJs) was added using a micropipette to each plastic box and the solution was allowed to spread evenly on the filter paper. Five 5^th^ instar larvae were collected using fine insect handling forceps and were shifted into each box containing nematodes.

The plastic boxes were then sealed and incubated for four days at 20°C. After the incubation period, the plastic boxes were carefully examined to record data on the mortality of weevil’s larvae. The larvae infected with nematodes were then dissected under the stereomicroscope in the ringer’s solution. The larvae were carefully observed for the presence of nematode in the body of dead larvae. The infestation percentage was calculated for statistical analysis.

### Infective potential of EPN’s against weevils’ larvae based on adult emergence

For the second experiment, the procedure described in 3.6.1.1. was followed and transparent plastic boxes (9 × 5 × 5) were used having Whatman No. 1 filter paper in the bottom of the boxes and 100 IJs in 0.1 ml water were added. Five uniform age last instar larvae of red palm weevil were introduced in each plastic box. The experiment was replicated five times for each nematode specie. The plastic boxes were sealed using Para film to inhibit the moisture loss. The plastic boxes were incubated at 20°C until the last adult emerged. The data were noted on number of adults emerged and nematode species and converted into percent emergence.

### Infective potential of EPN’s against red palm weevil pupae based on adult emergence

The pathogenicity of four selected nematode species *i.e., H. Becterophora, S. faltiae, S. carpocapsae* and *S. glesri* against red palm weevil pupae was evaluated following the procedure described in 3.6.1.1 and 3.6.1.2 with the difference that instead of treating larvae, the newly formed pupae of red palm weevil were treated with the selected nematode species. For the investigations, pupae formed within the past 24 h were used. Five pupae per treatment were added to each plastic box having treated filter paper at the bottom of each box. The pupae retained in the treated boxes until the emergence of the adult weevils.

The damaged pupae were then dissected under the stereomicroscope in the ringer’s solution and were carefully checked to find out the nematode in the body of dead pupae to ensure the cause of the death of pupae.

### Field Studies

The experiments were conducted at research area of the department of Entomology, Gomal University, Dera Ismail Khan to investigate the infectivity/pathogenicity of four different species of entomopathogenic nematodes *i.e., H. Becterophora, S. faltiae, S. carpocapsae* and *S. glesri* against red palm weevil.

For this purpose, the date palm plants of age 2 years were used. The date palm plants were grown under lath house conditions to prevent them from the insect infestation. The experiment was laid out as randomized complete block design having five repeats each consisting of five treatments including control. For each treatment five date palm plants were selected. The date palms were artificially infested with 5 larvae of 10 days age early in the morning at 9am. For the release of larvae, a hole of 4cm deep and 1cm wide was drilled in the date palm trunk at 1m from the ground level by the help of electric drill machine slanting down at 45° angle. The larvae were released in the trunk and the hole of the trunk was plugged with cotton covered with mud to inhibit the escape or entry of new weevils. Similarly, after 5 days of the release of weevil’s larvae, another hole of same size was drilled to the opposite side of the previous hole in the trunk and infested with 5 larvae of 15 days old age and the hole was concealed after the release of larvae. After 30 days of 1st release of the larvae, the palm trunks were injected with 6000 IJs nematodes. The infective capacity of selected four strains of nematodes was investigated. There were five treatments comprising of *H. bacterophora* (6000)*, S. carpocapsae* (6000)*, S. faltiae* (6000), *S. glesri* (6000) and control. The palm trunks were dissected 2 weeks after the application of 1^st^ (IJs) nematodes suspension. The data on the number of dead red palm weevil were recorded. To conform, the cause of weevil’s mortality, all the dead weevils were dissected and were carefully observed under microscope. The data was noted on the mortality of weevils and were converted to % mortality.

### Statistical analysis

The documented data were analyzed statistically using one-way analysis of variance technique and means were separated using Least Significance Difference (LSD) test using α = 0.05. The statistical analysis was carried out using computer software (SPSS ver. 13).

## Results

### Pathogenicity of EPN’s against Red palm weevil larvae based on dissection

The percent infestation of larvae based on dissection was found statistically higher in all the nematode species treated larvae compared to control. Among the different species of nematodes, the *S. carpocapsae* was found most effective with 94.68% nematode infested larvae followed by *H. bectrophora* with 92.68% infestation having significant variation between each other. The minimum number of 67.60 and 70.88% nematode infested larvae were found in *S. glesri* and *S. feltiae* treated larvae. Similarly, all the tested nematode species had significant effect on the percent adult emergence of red palm weevil. Among the nematode species, the minimum adult emergence (4.60%) of weevil was documented in *S. carpocapsae* treated larvae showing the significantly higher pathogenicity compared to other tested nematode species. The maximum adult emergence was noted in *S. glesri* treated larvae. Overall, maximum (97.80%) adult emergence was noted in untreated weevils.

### Infective potential of EPNs against red palm weevil pupae based on adult emergence

All the tested nematode species had significantly higher pathogenicity against pupae of red palm weevil compared to control. The *S. carpocapsae* was found to be the most effective species with 83.60% infested pupae having significant variation from all the tested nematode species. The *S. glesri* and *S. feltiae* were found least pathogenic nematodes having 43.60% and 44.80% infested pupae. No infested pupae were found in control/untreated treatment. As for as the adult emergence in concerned, the maximum adult emergence was noted in *S. glesri* treated pupae having non-significant variation from *S. feltiae* treated pupae. Among the treatments, the minimum adult emergence of 15.68% and 19.56% was noted in *S. carpocapsae* and *H. bectrophora* treated pupae. Overall, the maximum adult emergence of 98.40% was documented in untreated/control treatment.

### Effect of EPNs on the mortality of 6th instar larvae

The data presented in Fig. 4.6.1 indicate that all the tested nematode species were found highly pathogenic against 6^th^ instar larvae of red palm weevil. The mortality of the weevils gradually increased with an increase in the exposure period with maximum mortality recorded after 340 hours of the treatment. The first mortality of the treated larvae was noted at 24 hours after exposure period only in *S. carpocapsae* treated larvae, whereas; *H. bectrophora* caused first mortality after 48 hours and *S. feltiae* and *S. glesri* required 72 hours to produce first mortality of weevils. Among the tested nematode species, the *S. carpocapsae* was found most effective compared to other evaluated nematode species, with 100% mortality at 340 hours after treatment. It was followed by *H. becterophora* with 96% mortality. The *S. glesri* and *S. feltiae* were found least pathogenic nematode species registering 70 and 76% mortality of 6^th^ instar red palm weevils’ larvae at 340 hours after treatment. No mortality of red palm weevil larvae was documented during 10-days/340 hours exposure period in control treatment.

**Fig 1.**
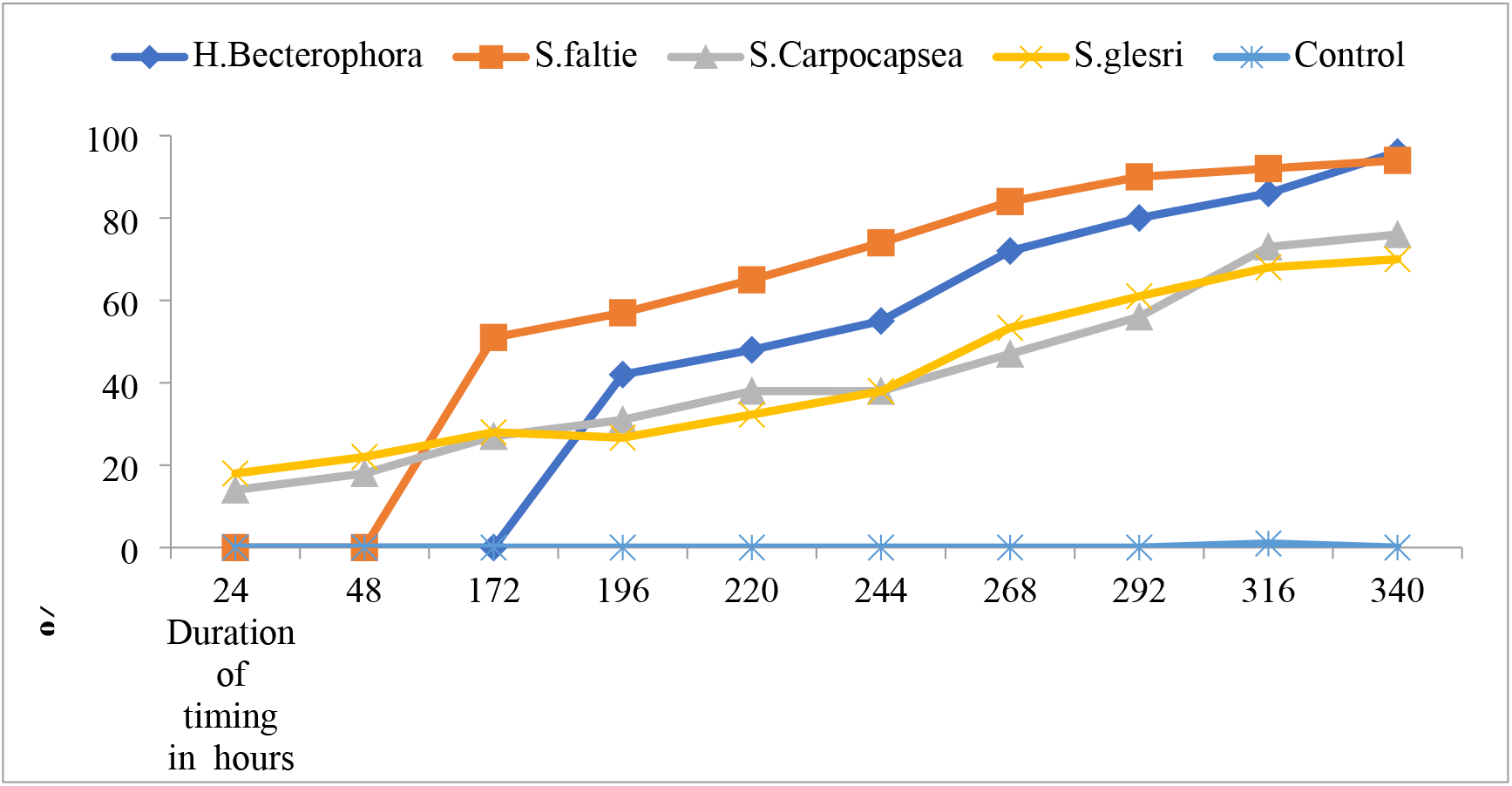
Effect of various species of entomopathogenic nematodes on the mortality of 6^th^ instar larvae of red palm weevil.

### Infective potential of EPN’s against red palm weevil adults

All the four evaluated species of nematodes were found highly pathogenic against the adult weevils (Fig. 4.6.2). Among the tested nematodes, the *S. carpocapsae* were found most pathogenic followed by *H. bectrophora*. No mortality of adult weevils was observed in first three days after the treatment of adult weevils. The first mortality of adults was recorded at four days after the exposure period in *S. carpocapsae* treated adults. The *H. bectrophora* required five days to show pathogenic effects against adult weevils. The *S. feltiae* and *S. glesri* were found least pathogenic and required six days to cause first mortality in adult weevils. The maximum mortality of 70% was observed in adults after 16 days of the treatment in *S. carpocapsae* treated adults followed by 63% mortality in adults treated with *H. bectrophora*. The minimum mortality of 59 and 58% was observed in adults treated with *S. feltiae* and *S. glesri* at 16 days after treatment. No further mortality of adult weevils was observed from 16 to 18 days after treatment with nematodes.

**Fig 2.**
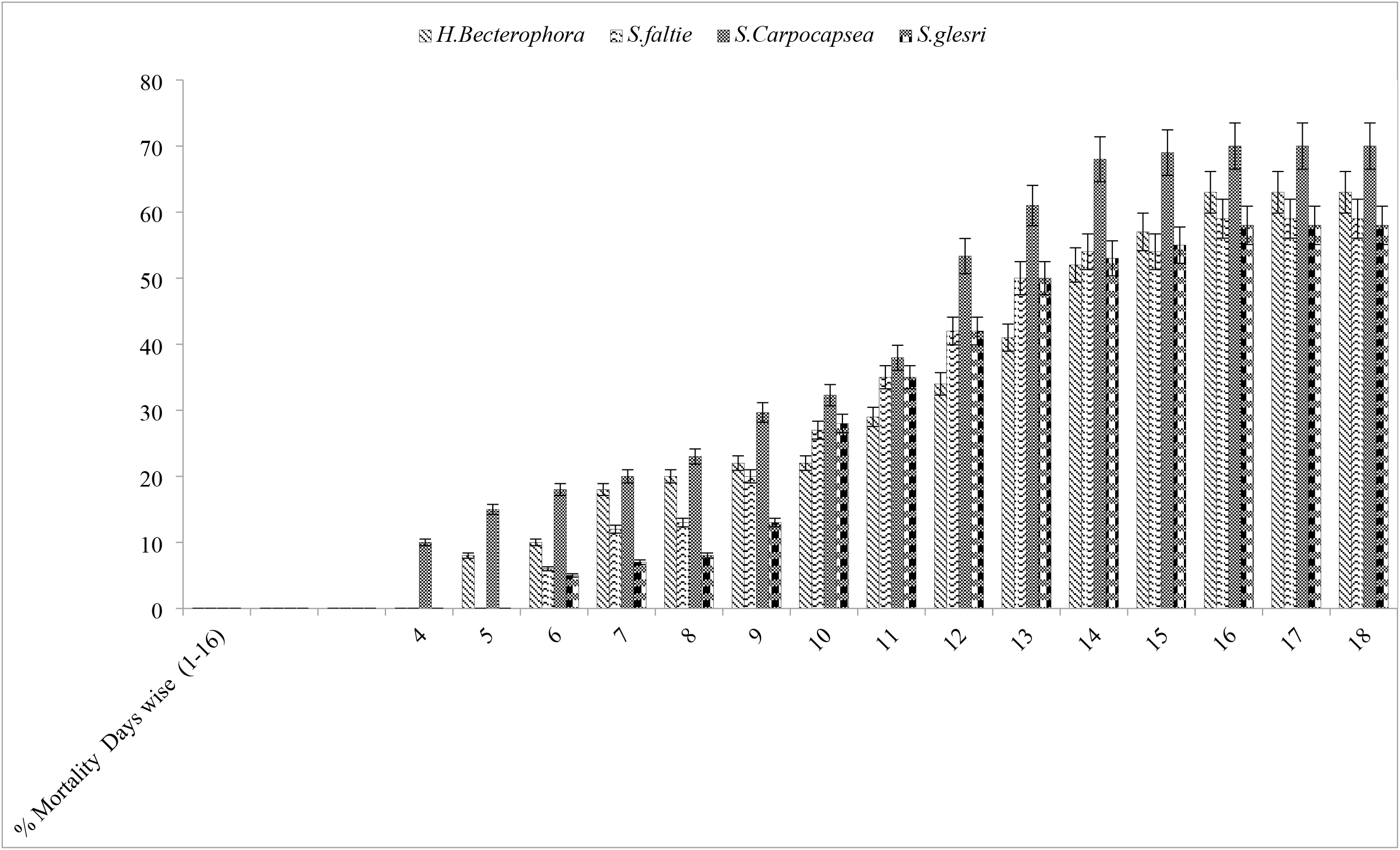
Effect of various species of entomopathogenic nematodes on the mortality of adult red palm weevil under laboratory conditions.

### Infective potential of EPN’s against red palm weevil under field conditions

All the tested nematode species were found equally pathogenic against the red palm weevil under field conditions. The *S. carpocapsae* were found maximum effective causing 83.60% mortality of red palm weevil. Among the tested four species of nematodes, the *S. glesri* and *S. feltiae* were found least effective with 60.60 and 64.80% mortality having significant variation from each other.

## Discussion

Biological control strategies can yield promising results without deteriorating the quality of the products by managing concealed insect pests of different tree species. The entomopathogenic nematodes play a pivotal role as bio-control agents of different species of insect pests including red pal weevil. They actively look for their target by utilizing host seeking ability and invade the target by entering the insect body through natural openings. In current investigations, the entomopathogenic nematodes belonging to Steinernematidae and Heterorhabditidae families were found highly pathogenic against larvae, pupae and adult stages of red palm weevil. In the present studies, among the tested different species of nematodes, the *S. carpocapsae* was found most effective with 94.68% nematode infested larvae followed by *H. bectrophora* with 92.68% infestation. The minimum number of nematode infested larvae were found in *S. glesri* and *S. feltiae* treated larvae. The larval infestation percentage (94.68%) of *S. carpocapsae* is lower than that reported in Italy [30]. They reported that the *S. carpocapsae* caused 100% mortality of red palm weevil larvae. Similarly, all the tested nematode species had significant effect on the percent adult emergence of red palm weevil. The minimum adult emergence was documented in *S. carpocapsae* treated larvae. The results are in agreement with [14]. They found that the maximum mortality (91.4%) of red palm larvae was recorded with *S. carpocapsae.* Comparable results were also reported by [21]. They documented that the nematode *S. carpocapsae* was found highly effective causing 98% mortality of red palm weevil under field conditions.

Similarly, all the tested nematode species had significant pathogenicity against pupae of red palm weevil compared to control. The *S. carpocapsae* was found to be the most effective species with 83.60% infested pupae whereas; the *S. glesri* and *S. feltiae* were found least pathogenic nematode strains. Similar result regarding pathogenicity of nematodes against red palm weevil were also recorded by [22]. They documented that among the three evaluated nematode species, the *S. carpocapsae* were found highly pathogenic against 3^rd^ and 10^th^ larval instar of red palm weevil causing 96.5% and 88.17% mortality. They also reported that the *S. feltiae* were found least effective causing only 38.68% and 35.35% mortality of 3^rd^ and 10^th^ instar larvae of red palm weevil. Similar findings were also documented by [35]. The *S. carpocapsae* was found most effective causing maximum mortality of 96.4% of last instar larvae of red palm weevil under laboratory conditions.

All the tested nematode species were found highly pathogenic and caused significant mortality in 6^th^ instar larvae of red palm weevil. The mortality of the weevils gradually increased with an increase in the exposure period with maximum mortality recorded after 340 hours of the treatment. The *S. carpocapsae* caused first mortality of the treated larvae at 24 hours after exposure period whereas; *H. bectrophora* required 48 hours to cause first mortality of weevils. The Sternematids have been reported more effective compared to heterorhabditids against red palm weevil larvae [29]. Overall; the nematodes were most effective against larvae (5^th^ and 6^th^ instar) followed by adult weevils, whereas; the tested species of nematodes were found least effective against pupae of red palm weevil. These results are supported by previous studies [36]

All the tested nematode species were found equally effective against the red palm weevil under field conditions. Among the tested nematodes, the *S. carpocapsae* were found maximum effective causing 83.60% mortality whereas; the *S. glesri* and *S. feltiae* were found least effective with 60.60 and 64.80% mortality. The pathogenic potential of *S. carpocapsae* was similar to that reported by [29]. The *S. carpocapsae* caused maximum mortality of 96.4% in the last instar larvae of red palm weevil. In previous studies, the entomopathogenic nematodes have been found most pathogenic against larvae of red palm weevil compared to pupae or adult weevils [45]. This trend of pathogenicity has also been documented against pine weevil, *Hylobius abietis* [41]. The variation in the susceptibility of different life stage, with larval stages most susceptible for infection by the nematodes has also been observed with other species of insects [42–44].

## Conclusion

The obtained results revealed that *S. carpocapsae* were found equally pathogenic compared to other tested nematode strains both under laboratory and field conditions against different developmental stages of red palm weevil. The larval stage of red palm weevil is the most susceptible stage for infection with nematodes. The use of *S. carpocapsae* is recommended for the timely management of red palm weevil under natural field conditions.

**Table 1.** Effect of various species of entomopathogenic nematodes on the % infestation and adult emergence of red palm weevil based on 5^th^ instar larvae treatment.

| **EPN Specie** | **% infestation** | **% adult emergence (Mean ± SE)** |
| --- | --- | --- |
| *H. Becterothora* | 92.68 ± 0.41 b | 6.60 ± 0.54 d |
| *S. feltiae* | 70.88 ± 0.29 c | 28.60 ± 0.54 c |
| *S. carpocapsae* | 94.68 ± 0.34 a | 4.60 ± 0.54 e |
| *S. glesri* | 67.60 ± 0.37 c | 32.40 ± 0.54 b |
| Control | 0.00 ± 0.54 d | 97.80 ± 1.13 a |
| LSD Value | 1.34 | 0.64 |

Mean ± standard deviation. Means in a column sharing similar letters are statistically similar at 5% level of probability using LSD test.

**Table 2.** Percent infestation and adult emergence based on treated pupae of red palm weevil.

| **EPN Species** | **% Infestation pupae** | **% adults emerged (Mean ± SE)** |
| --- | --- | --- |
| *H. Becterothora* | 80.20 ± 0.44 b | 19.56 ± 0.51 c |
| *S. feltiae* | 44.80 ± 0.83 c | 56.20 ± 0.83 b |
| *S. carpocapsae* | 83.60 ± 0.54 a | 15.68 ± 0.64 e |
| *S. glesri* | 43.60 ± 1.14 a | 56.40 ± 0.54 d |
| Control | 0.20 ± 0.44 d | 98.40 ± 0.51 a |
| LSD | 0.96 | 0.83 |

Mean ± standard deviation. Means in a column sharing similar letters are statistically similar at 5% level of probability using LSD test.

**Table 3.**
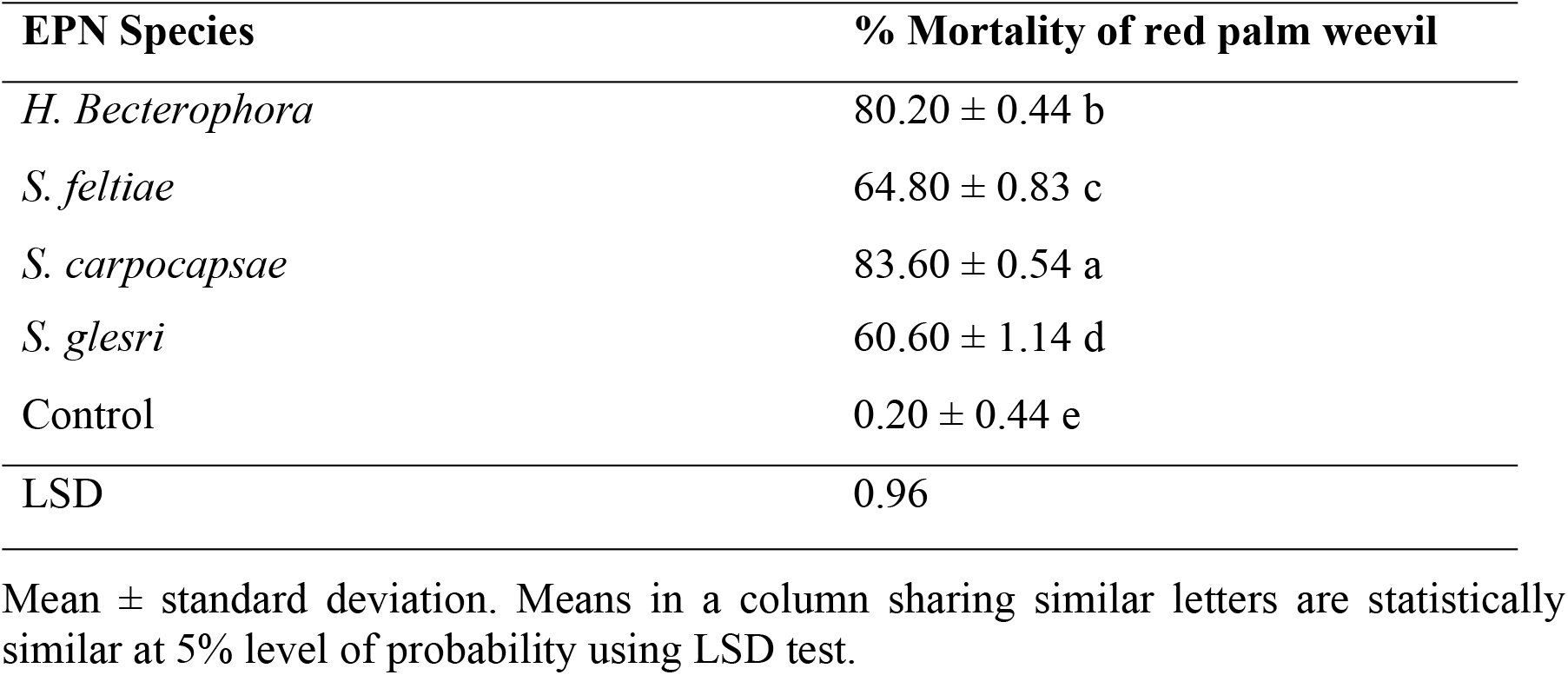
Effect of various species of entomopathogenic nematodes on the mortality of red palm weevil under field conditions.

| EPN Species | % Mortality of red palm weevil |
| --- | --- |
| *H. Bacterophora* | 80.20 ± 0.44 b |
| *S. feltiae* | 64.80 ± 0.83 c |
| *S. carpocapsae* | 83.60 ± 0.54 a |
| *S. glesri* | 60.60 ± 1.14 d |
| Control | 0.20 ± 0.44 e |
| LSD | 0.96 |

Mean ± standard deviation. Means in a column sharing similar letters are statistically similar at 5% level of probability using LSD test.

## Author Contributions

Gul Rehman conducted the experiments, Muhammad Mamoon-ur-Rashid Conceptualized, wrote the article and analyzed the data, Atiq Ahmad Alizai, helped in data collection, review and editing of this manuscript. This article is based on the Ph.D. research work of the principal author and has not been submitted to any journal except to Higher Education Commission, Pakistan.

